# Region-specific mean field models enhance simulations of local and global brain dynamics

**DOI:** 10.1101/2025.01.22.634225

**Authors:** Roberta Maria Lorenzi, Fulvia Palesi, Claudia Casellato, Claudia A.M. Gandini Wheeler-Kingshott, Egidio D’Angelo

**Author notes:** <u>corresponding author:</u>. **Author contribution:** ED and CGWK conceptualized the study. RL designed the pipeline and performed the analyses. RL, and ED wrote the paper. FP and CC provided support and guidance with data analysis. All the authors reviewed the submitted version of the manuscript.

## Abstract

Brain dynamics can be simulated using virtual brain models, in which a standard mathematical representation of oscillatory activity is usually adopted for all cortical and subcortical regions. However, some brain regions have specific microcircuit properties that are not recapitulated by standard oscillators. Moreover, magnetic resonance imaging (MRI)-based connectomes may not be able to capture local circuit connectivity. Region-specific models incorporating computational properties of local neurons and microcircuits have recently been generated using the mean field (MF) approach and proposed to impact large-scale brain dynamics. Here we have used a MF of the cerebellar cortex to generate a mesoscopic model of the whole cerebellum featuring a prewired connectivity of multiple cerebellar cortical areas with deep cerebellar nuclei. This multi-node cerebellar mean field model was then used to substitute the corresponding standard oscillators and build up a cerebellar mean field virtual brain (cMF-TVB), for a group of healthy human subjects. Simulations revealed that electrophysiological and fMRI signals generated by the cMF-TVB significantly improved the fitness of local and global dynamics with respect to a homogeneous model made solely of standard oscillators. The cMF-TVB reproduced the rhythmic oscillations and coherence typical of the cerebellar circuit and allowed to correlate electrophysiological and functional MRI signals to specific neuronal populations. In aggregate, region-specific models based on MF technology and pre-wired circuit connectivity can significantly improve virtual brain simulations fostering the generation of effective brain digital twins that could be used for physiological studies and clinical applications.

## 1. Introduction

Structure, function, and dynamics, the organizing principles dominating brain activity^1^, have recently been recapitulated by the virtual brain technology^2–4^. Virtual brains leverage structural Magnetic Resonance Imaging (MRI) data to reconstruct brain connectivity, place mathematical representations of neural activity in grey matter nodes, and generate emerging dynamics through simulations^5–7^ In this way, microscale neuronal dynamics can be integrated into mesoscale population models, which act then as generative models for macroscale signals such as Blood Oxygen Dependent (BOLD) signals, acquired with functional MRI (fMRI), or neural activity generated by electrical currents recorded with electroencephalography (EEG) or magnetoencephalography (MEG)^3,8,9^. Since virtual brain models can be tailored to single subjects, they are opening the perspective of brain digital twins^10,11^ applicable to personalized medicine for understanding brain function and enabling the personalized treatment of diseases.

The resolution of local microcircuit structure and function is critical to accurately simulate brain dynamics and extract physiological information at higher scales. While, ideally, the best resolution should be achieved for each node, in practice the mathematical representation of nodal activity has been simplified in previous modelling attempts. “The Virtual Brain” (TVB), the neuroinformatic platform commonly used to simulate virtual brain models^2–4,11–17^, operates under the assumption that a neural mass model, collapsing neuronal heterogeneity into two populations, (one excitatory and the other inhibitory) can capture local microcircuit dynamics all over the brain^18,19^. Typical neural mass models include the Wong-Wang (WW), the Wilson-Cowan (WC), and the Generic Oscillator (GO) model^19–22^. However, by using the same simplified neural mass models for all nodes, TVB misses the opportunity to capture local microcircuit functional diversity. In contrast to neural mass models, *mean field* (MF) models maintaining the physiological structural organization and functional properties of local neuronal microcircuits have been developed and applied to main cortical microcircuits^23–28^. In particular, the cerebellum is a large brain structure amounting for more than half of all brain neurons^18^, is tightly interconnected with the rest of the brain^29,30^, and has a feed-forward organization that is not represented by standard neural masses^31^. The cerebellar cortex features four major neuronal populations, i.e., Granule Cells (GrC), Golgi Cells (GoC), Molecular Layer Interneurons (MLI), and Purkinje Cells (PC), whose physiological function and connectivity are oversimplified (or overtly wrong in physiological terms) in neural mass models. Moreover, the cerebellar output from PCs is inhibitory on Deep Cerebellar Nuclei (DCN), while communication between TVB nodes normally uses excitatory synapses only. Finally, communication between cerebellar nodes cannot be fully resolved by Magnetic Resonance Imaging (MRI) tractography. All these issues can severely compromise the anatomo-physiological consistency of virtual brain simulations. The cerebellar mean field model (CRBL MF) is tailored on salient features of the cerebellar neuronal populations based on precise cellular-level reconstructions of the microcircuit^26,31,32^. Structural and functional parameters of the CRBL MF, such as the synaptic convergence and the quantal synaptic conductance, were extracted from a Spiking Neural Network (SNN) previously validated using experimental recordings^31–34^. For methodological consistency, the mean field theory used to develop the CRBL MF follows that of the MF of the cerebral cortex, relying on a transfer function formalism that allows to transfer the microscale parameters to a mesoscale domain ^23,24^. Specificity of the cerebellar cortex neuronal architecture was preserved, including recurrent excitation and inhibition amongst GrC, GoC, MLI, and PC ^18^. The population-specific transfer functions can return postsynaptic activity from presynaptic inputs, accounting for neuronal firing rates and membrane potential fluctuations.

Here, we developed a *multi-node CRBL MF* and integrated it into TVB generating a *cerebellar mean field TVB* (*cMF-TVB*) model. This required first to develop a multi-node CRBL MF assembling interconnected MF (one for each cerebellar node) with nodes of DCN. Then, the multi-node CRBL MF was connected to the rest of the brain, in which nodes are typically represented as standard neural masses. To maintain the physiological interregional communication, the connectivity between cerebellar cortical nodes and DCN was imposed *a priori*, as it was not fully resolved on the microscopic and mesoscopic level by MRI tractography: inhibitory connections were set between PCs and DCN, 10% feedback was introduced from DCN to PCs, and parallel fibres (pfs) were added to connect adjacent cerebellar regions^35,36^. Finally, simulated electrical activity and the BOLD signal were referred to the specific MF neuronal populations most likely to generate them and were validated against empirical data recorded in humans *in vivo*. The cMF-TVB was validated against standard TVB simulations, constructed by associating the same two-population model to all the nodes. In aggregate, the cMF-TVB incorporating the multi-node CRBL MF proved capable to improve local circuit and whole brain simulations and to capture high-level physiological features of cerebellar activity like large-scale oscillations on multiple frequency bands^37,38^.

## 2. Results

In this work, we address how to integrate different types of models with MRI data into TVB platform using the CRBL MF as a prototype. Our framework allows us to address brain modeling using a multiscale approach which links the microscale neuronal circuitry information to the macroscale observed dynamics (Fig. 1). The technical implementation of the multi-node CRBL MF and its integration into TVB, ending up with the cMF-TVB, are detailed in Methods section.

**Fig. 1.**
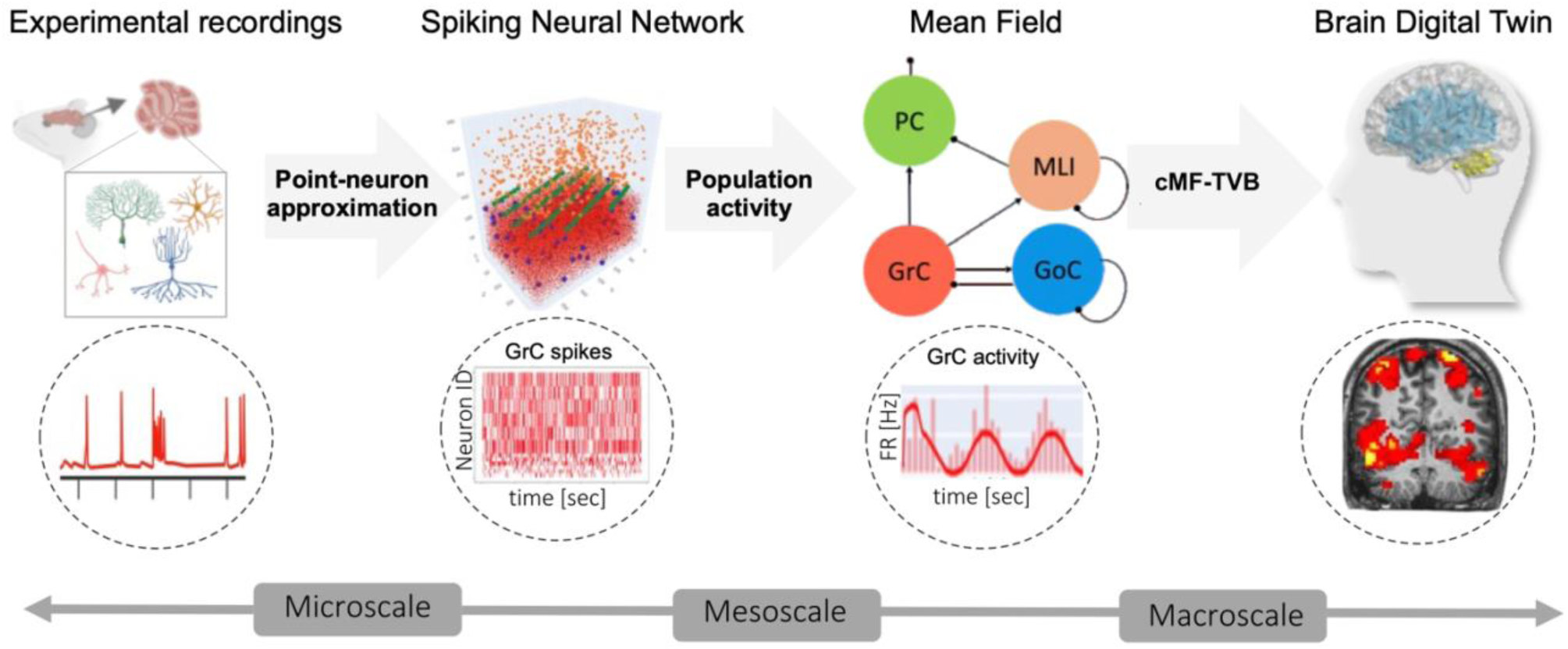
Multiscale brain modeling. An integrative approach to model cerebellar activity, bridging the gap between microscale and macroscale signals. Microscale experimental recordings (e.g., action potentials recorded in the GrC are used to develop point-neuron models). These are wired into highly specific microcircuits, e.g., a cerebellar spiking neural network reproducing the spiking activity of cerebellar neurons, such as GrCs. The spiking neural network serves as the backbone of the CRBL MF scaling up to the mesoscale by reproducing population activity in terms of firing rate [Hz]. The cerebellar mean field model is then integrated into a virtual-brain simulator, ending up with the cMF-TVB that provides effective multiscale simulations.

The nature of the CRBL MF, which is made of 4 neuronal populations required a specific curation of the structural connectivity (SC) to account for cerebellar-specific connectivity in the construction of the cMF-TVB circuitry, resulting in a curated cerebellar SC which includes the parallel fibers weights along with, the inhibitory projections from PC to the DCN (Fig. 2). The cMF-TVB is constructed by connecting the multi-node CRBL MF with the WW representing the DCN and the cerebrum, resulting in the circuitry in Fig3., which forms the functional back-bone of the cMF-TVB.

**Fig. 2.**
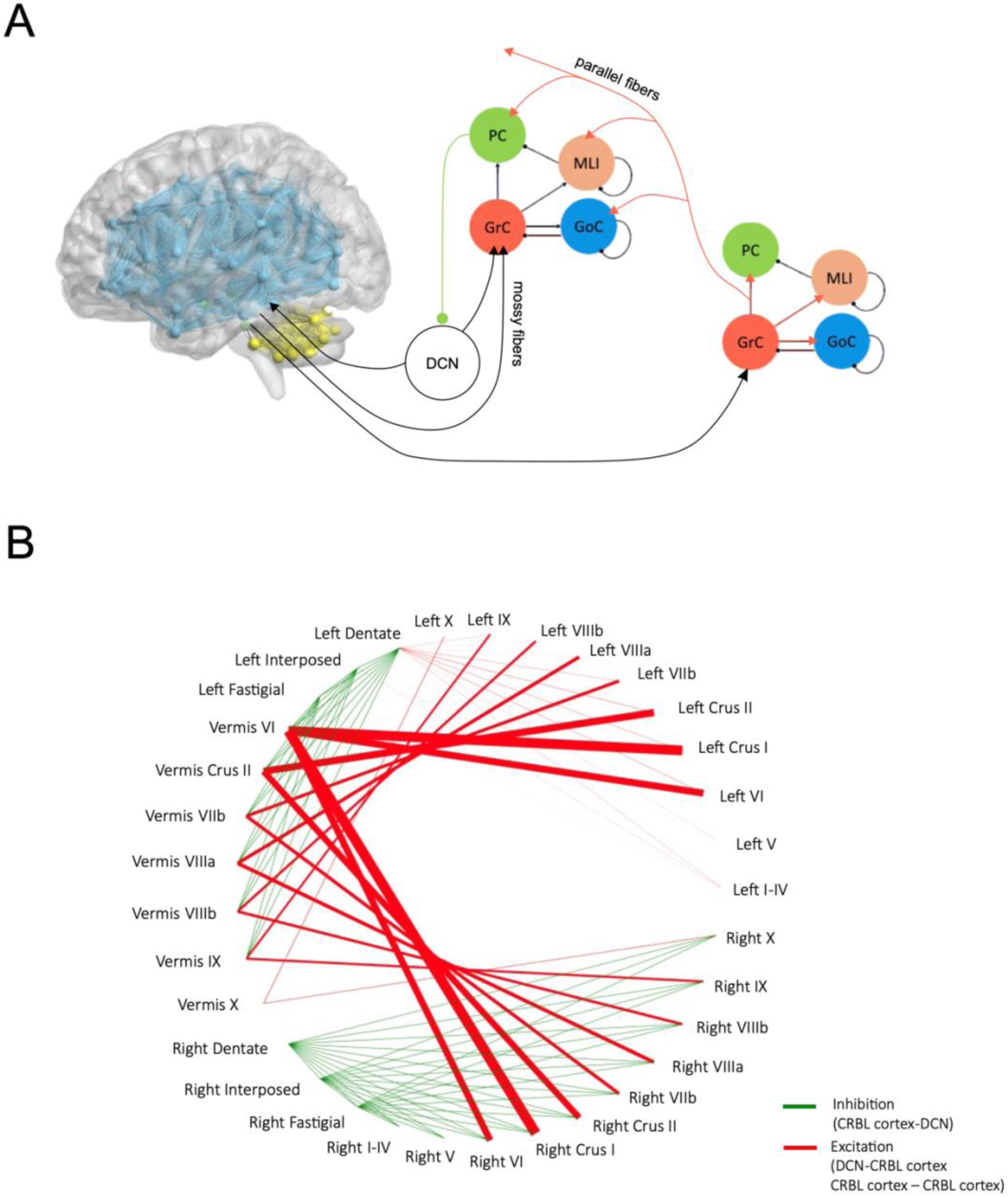
Curation of intra-cerebellar SC. **(A)** Cerebellar SC is curated by integrating microscale data with macroscale recordings. To quantify the connections between pairs of cerebellar cortical nodes, the convergence of parallel fibres from GrC to GoC, MLI, and PC are extracted from a cerebellar spiking neural network. Volumes of adjacent cerebellar regions are computed from subject-specific T13D images and summed to weight the population-specific synaptic convergences (i.e., GrC-GoC, GrC-MLI, and GrC-PC). Volume-weighted synaptic convergences are normalized for the Intracranial Volume. The connectivity between cerebellar cortex and DCN is derived from anatomically constrained tractography computed on the pre-processed subject-specific diffusion weighted images. The connection weights from cerebellar cortical nodes to the DCN nodes is made inhibitory and the feedback from DCN to cerebellar cortical nodes was reduced at 10% of the feedforward, accordingly to the physiology of the cerebellar circuit. **(B)** Curated cerebellar SC. Green edges represent inhibitory connections (from PC to DCN), while red edges represent excitatory connections made by parallel fibres between adjacent cerebellar cortex nodes. The thickness of the edges is proportional to the SC weights.

**Fig. 3.**
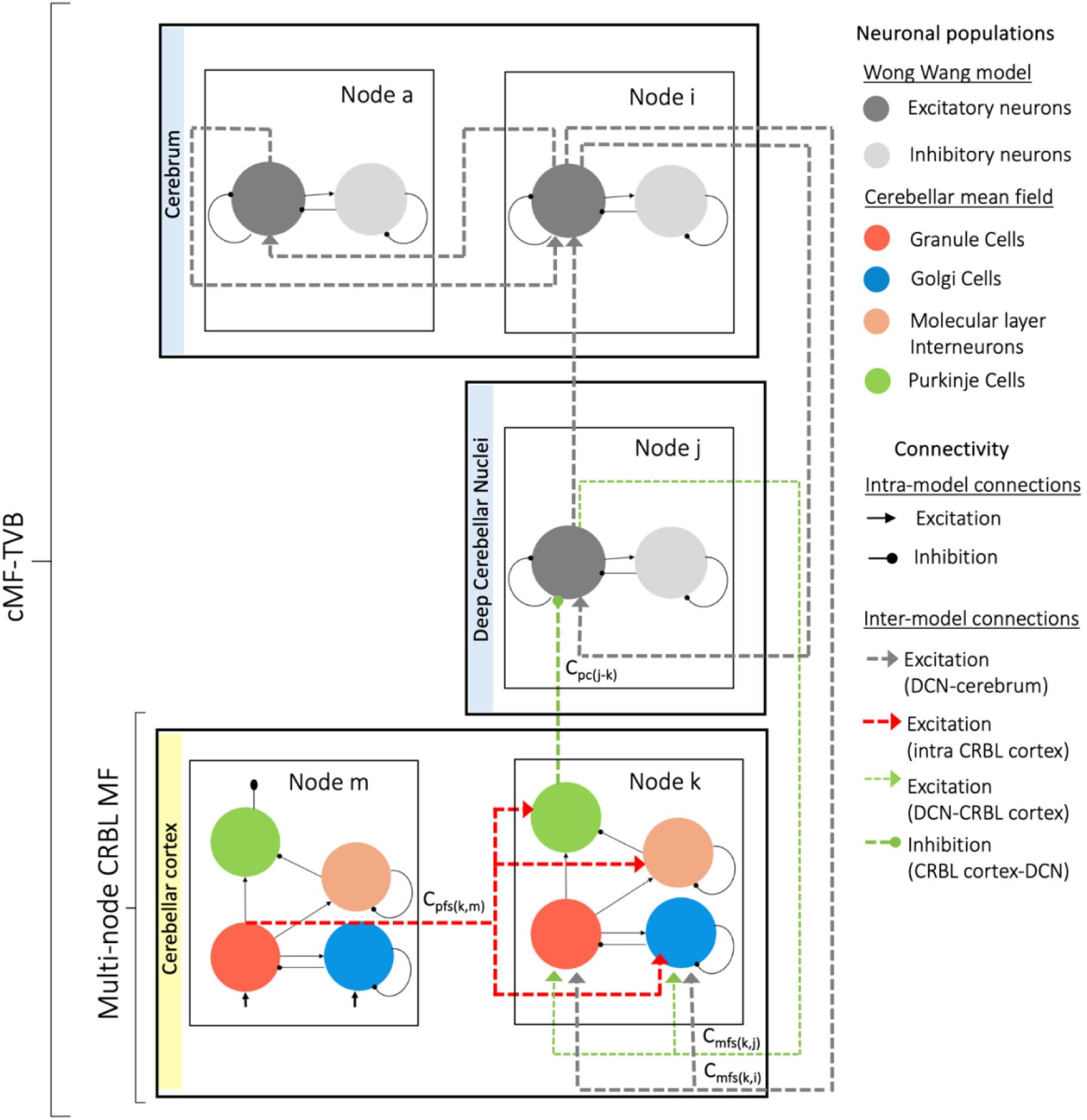
Schematic circuit representation of the cMF-TVB. The cMF-TVB is built by interconnecting different models according to the spatial map defined by the SC. The cMF-TVB includes the multi-node CRBL MF, made of 27 MF models in the cerebellar cortex, hardwired with 6 WW models in DCN (yellow box), plus 93 WW models in the other brain nodes (light blue boxes). After segmentation and remapping in an atlas virtual space, each cerebellar region is attributed to a node and is represented with a MF. Connectivity between adjacent CRBL MFs laying on the transverse plane corresponds to parallel fibres (pfs) generated by GrCs in a source node (m) projecting its activity to GoC, MLI and PC in a target node (*k*), so that the coupling strength [Hz] from *m* to *k* through parallel fibers (pf) is *C_pfs(k,m)_*. Node *k* forwards the inhibitory activity of its PCs to the excitatory neurons of a DCN (*j*), which then sends back connections to node k, with coupling strength *C_PC(j,k)_*. and *C_mfs(k,j)_*, respectively. Node k also receives the excitatory activity of a cerebral node (*i*), so that the coupling strength [Hz] from *j* to *k* through mossy fibres (mf) is *C_mfs(k,i)_*. DCN and the rest of the brain are coupled through the activity of their excitatory neurons (coupling C between standard oscillators are omitted for simplicity).

Simulations of cerebellar dynamics were used to assess the constructive and predictive validity, specifically (i) to assess the robustness of the multi-node CRBL MF and CMF-TVB in simulating neuronal activity; (ii) to investigate the similarity between empirical BOLD (empBOLD) and simulated BOLD (simBOLD) computed by convolving the multi-node CRBL MF and cMF-TVB responses with the TVB built-in hemodynamic response function; and iii) to determine the ability of the cMF-TVB in capturing the multiple oscillatory bands of the cerebellum^37^.

### 2.1. Constructive and predictive validity

Simulations of cerebellar neuronal activity were performed both in open loop, i.e., isolated cerebellar cortex, represented by the multi-node CRBL MF (Fig. 4) and in closed loop, considering the cerebellar cortex connected to the DCN and the cerebrum, represented by the cMF-TVB (Fig5). The CRBL MF showed population activity within physiological ranges for GrC, GoC, MLI and PC, demonstrating that the biologically grounded features of a single CRBL MF^26^ was preserved after the multi-node integration in TVB. Simulations of the BOLD signal were performed both for the multi-node CRBL MF (open loop) and the cMF-TVB (closed loop). The performance of the cMF-TVB was assessed either representing the CRBL with standard neural mass models or with the CRBL MF model and comparing the simBOLD with the empBOLD. Importantly, there is evidence that the BOLD signal of the cerebellum mostly reflects the on-off activity of neurons at the input stage (PC activity is almost stationary over the integration time length of fMRI)^39,40^, thus we specifically computed the simBOLD signal from GrC activity.

**Fig. 4.**
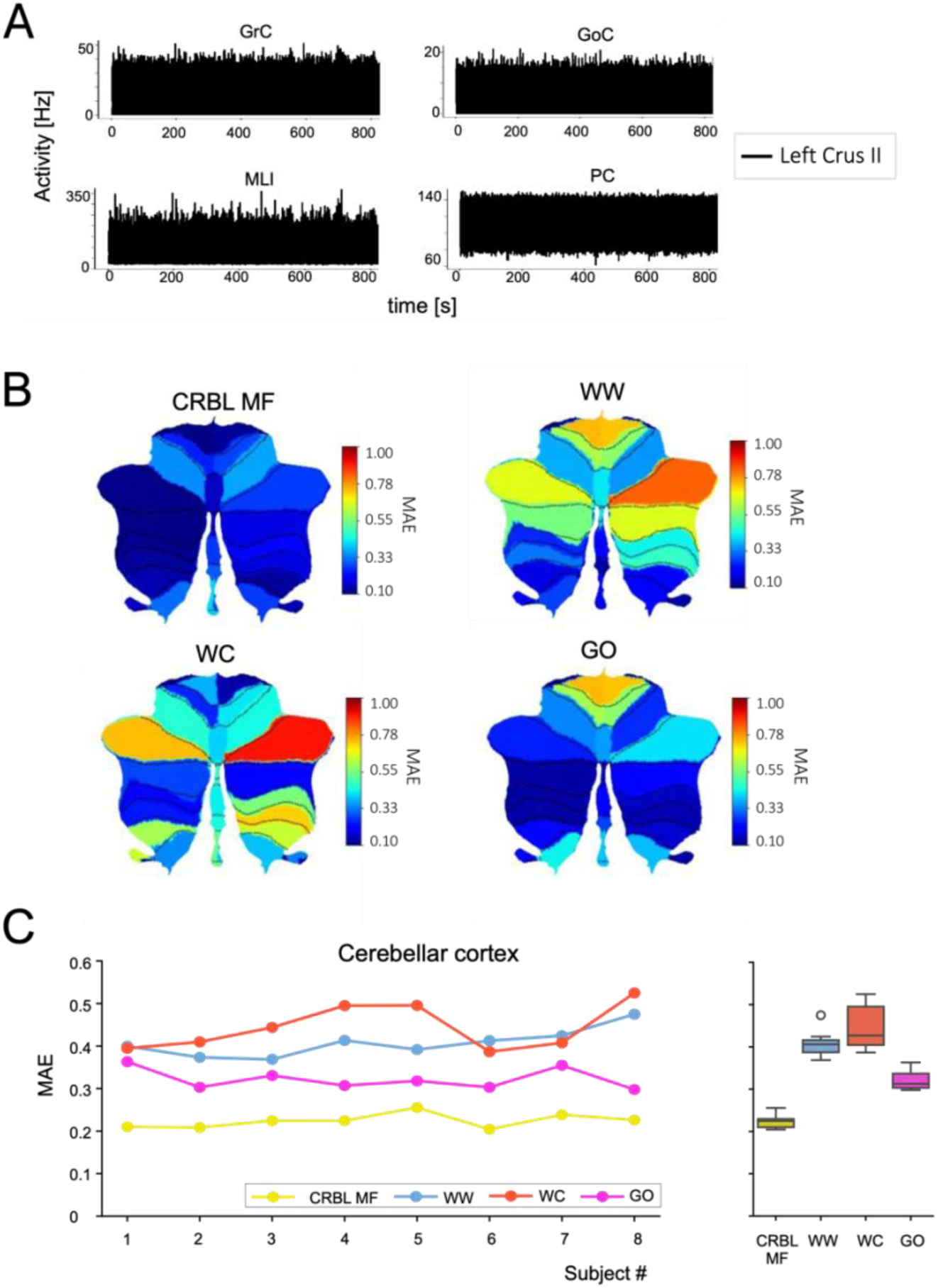
Multi-node CRBL MF. **A)** Cerebellar cortex neuronal activity, reported for left crus II as an example, is simulated using the CRBL MF integrated in the TVB platform for a randomly chosen subject taken from the Human Connectome Project dataset. The curated cerebellar structural connectivity was used to set the connections between different nodes (i.e., regions). To simulate the cerebellar cortex neuronal activity, one CRBL MF was associated with each node belonging to the cerebellar cortex. For each node, the simulated activity lays within the physiological range for each neuronal population. **B)** BOLD signals are extracted from fMRI data (emBOLD) and simulated using, as generative model, either the CRBL-MF or one of the generic unspecific models already available in the TVB platform (WW = Wong Wang, WC = Wilson Cowan, GO = Generic Oscillator). The flat maps show the MAE between empirical and simulated BOLD, averaged across subjects for each cerebellar cortex region (cold-colors correspond to a low MAE). The CRBL MF, as generative model for BOLD signals, remarkably reduces the MAE in each cerebellar cortex region. **C)** The single subject MAE is computed for each subject by averaging the region-specific MAE over regions. When using the CRBL MF as generative model for simulating BOLD signals, the MAE between simulated and empirical signals (∼0.20) is significantly reduced compared to unspecific models (Mann Whitney p-value <0.001)

**Fig. 5.**
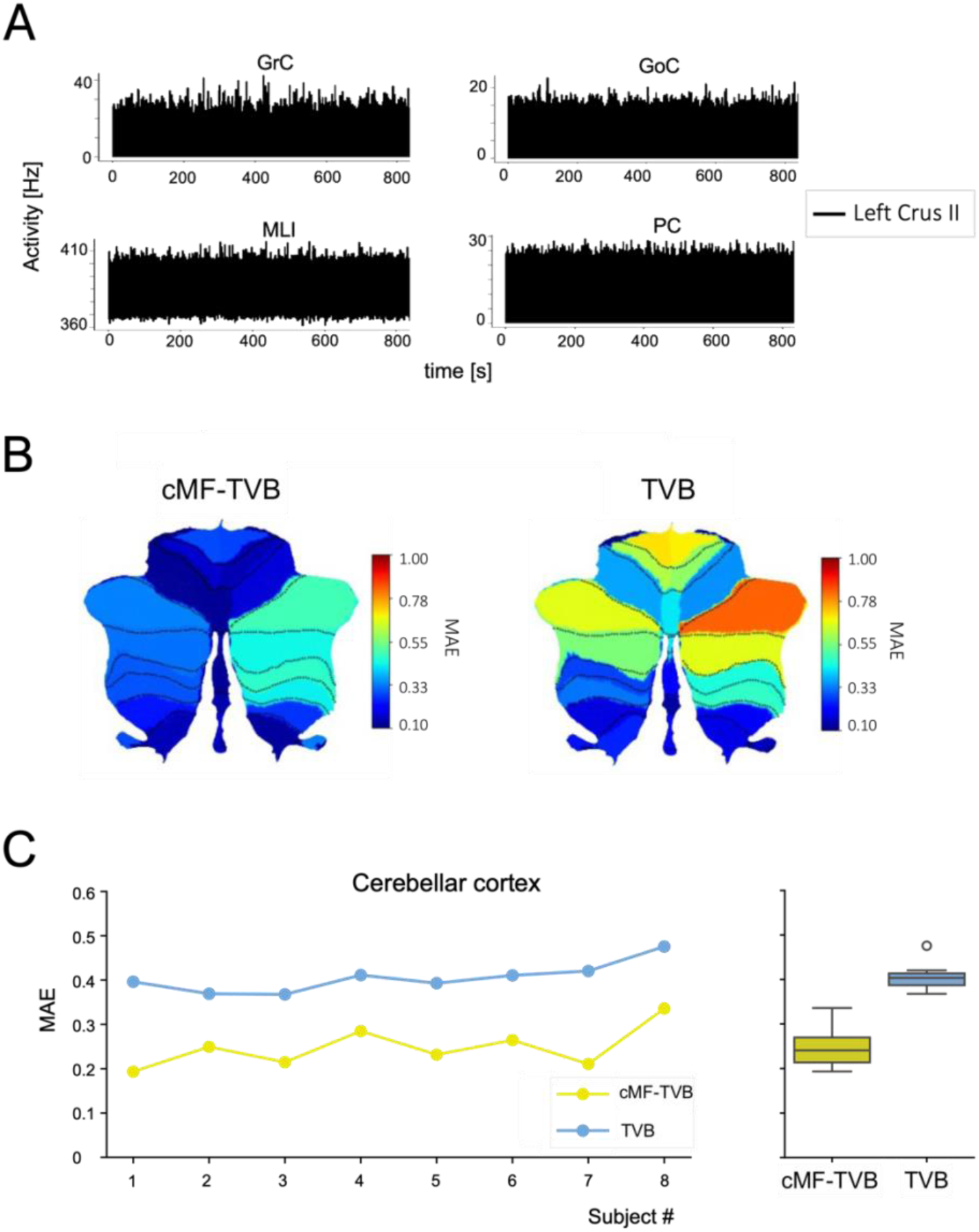
cMF-TVB simulation of the cerebellar cortex activity. Cerebellar cortex dynamics are evaluated from whole brain dynamics simulated with cMF-TVB and compared to a simulation performed using the standard TVB. **A)** Cerebellar cortex neuronal activity, reported for left crus II as an example, is simulated using the CRBL MF integrated in the TVB platform for a subject randomly chosen from the Human Connectome Project dataset. Curated whole brain structural connectivity is used to map the connections between whole brain nodes. To simulate the cerebellar cortex neuronal activity, a CRBL MF is associated with each node belonging to the cerebellar cortex, while WW is used for cerebral cortical and subcortical nodes. For each node, the simulated activity is within the physiological ranges for each neuronal population. Furthermore, the interconnection with cerebral nodes results in a heterogenous activity of MLIs and PCs, driven by the heterogeneous weight input form the cerebral regions to the GrCs. **B)** The flat maps show the MAE between empirical BOLD signals (extracted from fMRI data) and the simulated BOLD averaged over the subjects for each cerebellar cortex regions (cold color correspond to low MAE) extracted from the whole brain simulation. cMF-TVB shows a better performance compared to the standard TVB (WW models in all nodes), reducing the MAE in each cerebellar cortex region. **C)** The single subject MAE is computed for each subject by averaging the cerebellar cortex region-specific MAE over cerebellar cortex regions. When using the cMF-TVB for simulating whole brain activity, the MAE between simulated and empirical signals (∼0.25) is significantly reduced compared to the standard TVB (Mann Whitney p-value <0.001).

#### 2.1.1 Multi-node CRBL MF simulations (open loop)

In open loop simulations, the multi-node CRBL MF was activated by 4 Hz random activity on mossy fibres (mfs) of all the nodes. For each subject, the MAE between simBOLD and empBOLD was reduced when the multi-node CRBL MF was used instead of WW, WC, or GO (Fig. 4A). The same occurred with the average MAE across subjects (Fig. 4B). The Mean Absolute Error (MAE) was not-normally distributed for WW, WC, and GO, with a non-homogeneous variance in each model (Table 1). The MAE of the CRBL MF against other models was statistically different (Mann-Whitney test, p < 0.0001) for each pair of comparisons (Fig. 4C), namely: CRBL MF vs WW; CRBL MF vs WC; CRBL MF vs GO (Table 2).

**Table 1.**
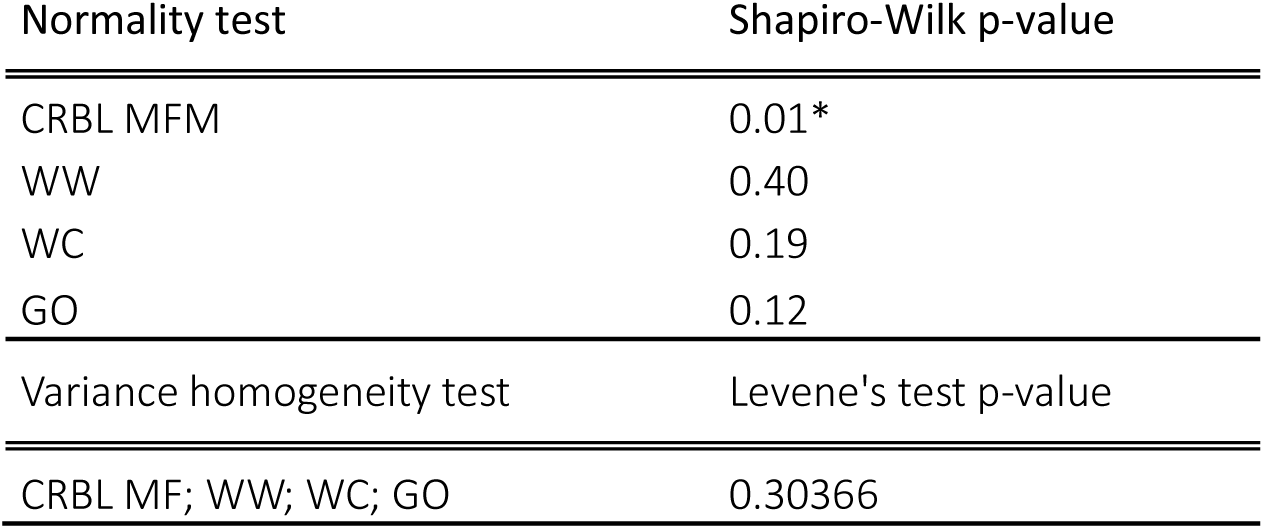
Analysis of the MAE distribution. A statistically significant threshold to reject the null hypothesis was set to 0.05 both for the Shapiro-Wilk and the Levene’s test. Statistically significant p-values (<0.05) are highlighted with *. Overall, MAE between empBOLD and simBOLD with the CRBL MF was not normally distributed (p-value < 0.05). The hypotheses of normality for the distribution of the MAEs between empBOLD and simBOLD generated by other models could not be rejected (p-value > = 0.05). Homogeneity of the variance across different models could not be rejected (p-value >= 0.05).

**Table 2.** Cerebellum open loop model comparisons. Comparisons between pairs of models (CRBL MF = cerebellar mean field, WW = Wong Wang, WC = Wilson Cowan, GO = Generic Oscillator) were performed using the Mann-Whitney test with statistically significant threshold set at 0.001. All comparisons resulted in statistically significant differences (p-value <0.001, marked with an asterisk) with the only exception of WW vs. WC.

| Models |  | Mann-Whitney p-value |
| --- | --- | --- |
| CRBL MF | vs. WW | * <0.0001 |
| CRBL MF | vs. WC | * <0.0001 |
| CRBL MF | vs. GO | * <0.0001 |
| WW | vs. WC | 0.2 |
| WW | vs. GO | * <0.0001 |
| WC | vs. GO | * <0.0001 |

#### 2.1.2 cMF-TVB simulations (closed loop)

In closed loop simulations the cerebellum was fed by activity transmitted by the other brain nodes of the cMF-TVB. The MAE between simBOLD and empBOLD, was calculated both for the TVB and cMF-TVB (Fig. 5 and Fig. 6). For each subject, the MAE between simBOLD and empBOLD in the cerebellar cortex, was reduced when the cMF-TVB was used instead of TVB (Fig. 5B). The same occurred with the average MAE across subjects (Fig. 5C). Interestingly, the MAE improvement was observed also for the DCN nodes (Mann-Whitney test, p < 0.001) (Fig. 6A) and for the whole brain (Mann-Whitney test, p < 0.001) (Fig. 6B), indicating that the improvement brought the CRBL MF propagated across the brain networks.

**Fig. 6.**
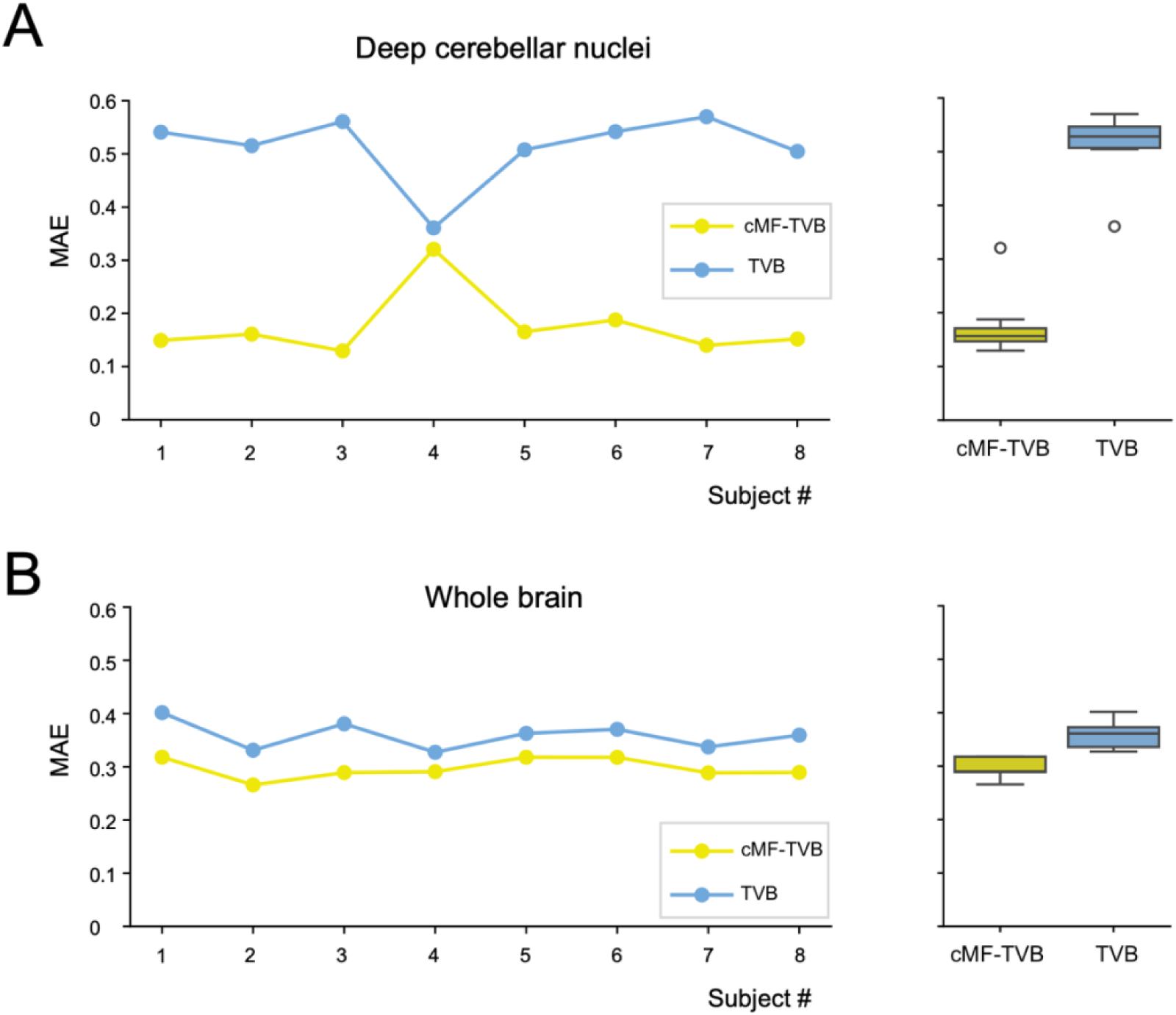
cMF-TVB simulation of the whole brain. Whole brain activity was simulated with cMF-TVB (one CRBL MF associated with each cerebellar cortical node and one WW associated with each deep cerebellar nucleus and cerebrum node). **A)** simBOLD signals of DCN were extracted from a whole brain simulations performed with either cMF-TVB or TVB. For each subject, simBOLD was averaged over regions and compared to empBOLD by computing the subject specific MAE for cMF-TVB and TVB. cMF-TVB improves DCN simulations for all the subject remarkably (except for subject 4, where cMF-TVB and TVB are comparable). Overall, the MAE is significantly reduced when using cMF-TVB (Mann Whitney p-value <0.001) showing that the inclusion of a region-specific model (i.e., CRBL MF) improves local dynamics compared to unspecific models (e.g., WW). **B)** The subject-specific MAEs between empBOLD and simBOLD using cMF-TVB and TVB are computed also for the whole brain simulation. For all the subjects, cMF-TVB improves the simulation performance significantly (Mann Whitney p-value <0.001), highlighting that region-specific models can improve also global dynamics.

### 2.2 Emergent cerebellar rhythms and coherence in the cMF-TVB

An emergent property of the cMF-TVB concerns its ability to reveal specific EEG/MEG bands in the cerebellar activity. The CRBL showed prominent activity in the theta band with relevant components also in other EEG bands (Fig. 7A). During resting-state cMF-TVB simulations, activity was recorded from all neuronal populations in the multi-node CRBL MF. Indeed, for each neuronal population, the Power Spectrum Density (PSD) showed activation in all bands (see supplementary material Fig. S3 for an example of a subject-specific PSD). At the group level, theta band activity predominated in GrC, MLI, and PC, while in GoC the delta, theta, and alpha bands had almost the same power (Fig. 7A). The predominant frequencies averaged across populations were 3 Hz for delta, 6 Hz for theta, 11 Hz for alpha, 20 Hz for beta, and 64 Hz for gamma (Fig. 7B). All bands were represented in the nodes of the multi-node CRBL MF, although some regional variations could be observed most probably reflecting the specific connectivity of each cerebellar node in the cMF-TVB and the different dynamics engaged in the local cerebellar circuits (Fig. 7C). For each region, population-specific carrier-frequency is reported in supplementary material (Table S1).

**Fig. 7.**
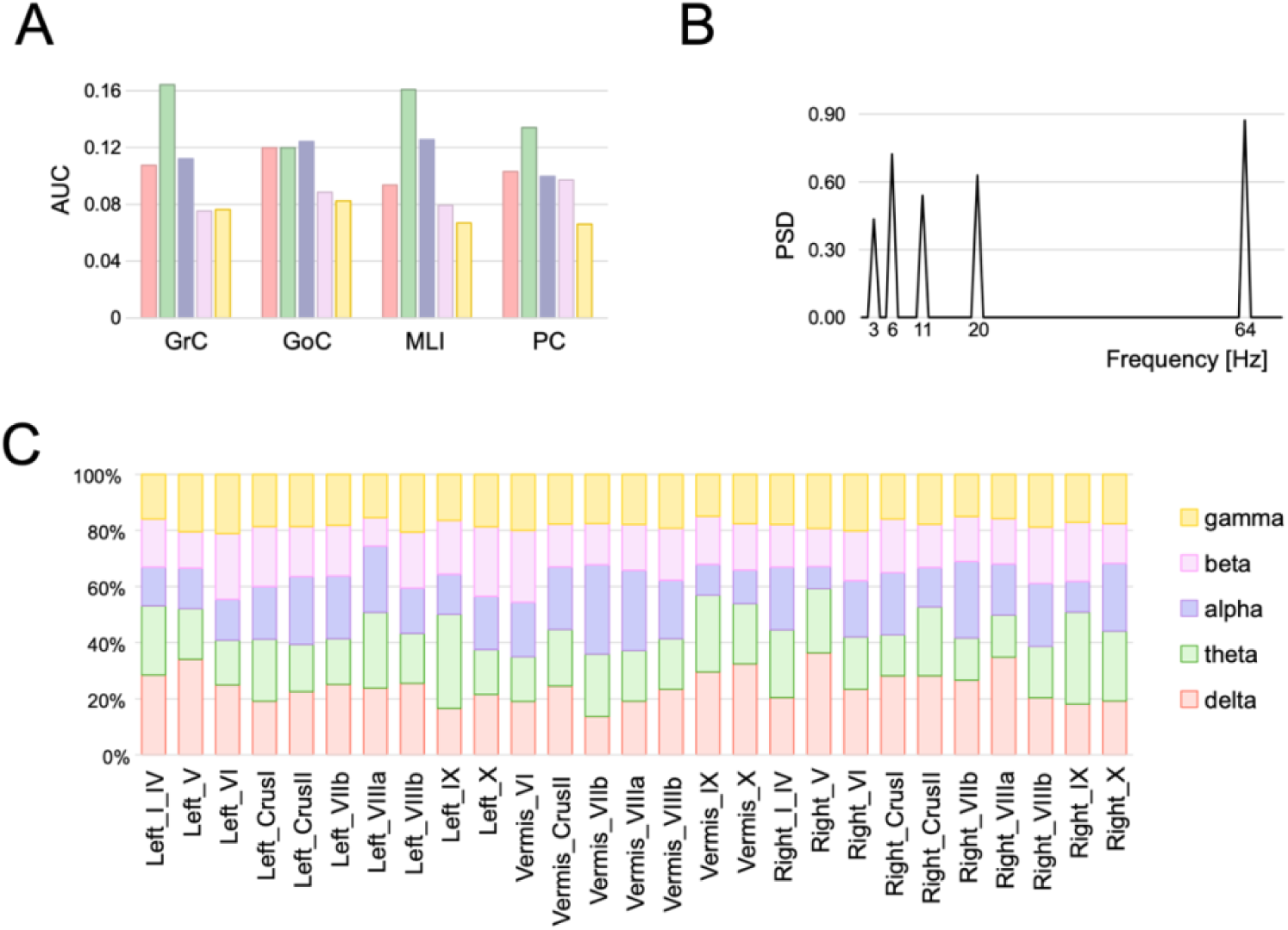
Cerebellar rhythms emerging from cMF-TVB simulations. **A)** For each population, the PSD is averaged across subjects to define the bands of circuit oscillations (delta = [0.4 4) Hz, theta = [4 8) Hz, alpha = [8 13) Hz, beta = [12 30) Hz, gamma = [30 100) Hz) for each neuronal population. The theta band is predominant in GrCs, MLIs, and PCs. **B)** Predominant frequencies for each band (only frequencies with a PSD >= 0.4 are reported). 64 Hz is the predominant frequency with a PSD = 0.87 revealing that a remarkable synchronization of cerebellar populations activity occurs in the middle of gamma band. **C)** Activity bands in different cerebellar regions. The weight of each band is computed by summing the population-specific power (i.e., the normalized PSD area under the curve, AUC). The gamma band is around 20% in all regions, the beta band is usually less than 20% (except in vermis VI and lobules X), the delta, theta, and alpha bands show higher variability across regions.

Another emergent property of the cMF-TVB concerns its ability to reveal coherence between cerebellar nodes during resting-state fMRI recordings. For all subjects, the coherence between cerebellar nodes was more similar to the empirical data when using the cMF-TVB instead of the TVB. The difference between the two models was statistically significant (see Frobenius norm, cosine similarity, Kullback-Leibler divergence and Pearson Correlation Coefficient in Table 3).

**Table 3.** Coherence of cerebellar subnetwork dynamics simulated with the cMF-TVB. cMF-TVB and TVB were compared separately with empirical data. Significant threshold of the t-test was set at 0.001 (MAE = Mean absolute error, KL = Kullback Leibler, PCC = Pearson Correlation Coefficient, SCC = Spearman correlation coefficient). The asterisk indicates statistically significant differences. Scores were selected to compare simulated coherence matrices with empirical coherence matrices under different perspectives: MAE to compute the magnitude of coherence, Frobenius Norm and cosine similarity to compare the discrepancy between matrices, KL divergence to evaluate the differences in distributions, and PCC together with SCC to investigate the coherence trend.

| Score | t-test p-value |
| --- | --- |
| MAE | 0.004 |
| Frobenius Norm | <0.001* |
| Cosine similarity | <0.001* |
| KL divergence | <0.001* |
| PCC | <0.001* |
| SCC | 0.600 |

## 3. Discussion

In the present work, for the first time, a multi-node CRBL MF model was developed and integrated into the cMF-TVB allowing to face a set of physiological issues that remained unaddressed so far. First, we have surpassed the generic representation of node activity in the TVB, accounting for the cerebellar microcircuit organization that does not conform to, e.g., WW, WC, or GC neural masses. Moreover, we have faced the inability of DWI to resolve fibres interconnecting cerebellar regions and DCN by pre-assembling the MF and DCN and by weighting their connectivity afterwards. Finally, according to physiology, we have modified the cerebellar output and made it inhibitory on DCN. It should be noted that the extra-cerebellar connectivity was also curated to obviate the inability of tractography in selecting out ipsilateral cerebello-thalamo-cortical spurious tracts^29,30,41^. We will consider below how the integration of anatomo-physiological properties of a specific brain circuit into large scale simulation advances the biomimetic capacity of virtual brains toward the generation of effective brain digital twins.

### 3.1 Architecture and function of the multi-node CRBL MF model

The CRBL MF has been designed to recapitulate the main anatomical and physiological features of the cerebellar cortex – DCN circuit and allow its simulation. The cerebellar cortex features a unique anisotropic geometry supporting a forward architecture with recurrent excitation and inhibition based on 4 main neuronal populations (GrC, GoC, MLI, and PC). These properties, along with the specific transfer function of neurons, have been incorporated into a MF model providing a biomimetic representation of a cerebellar region such as a node in a virtual brain^26^. Multiple such MF models have been interconnected to generate the multi-node CRBL MF model, which has been hard-wired with DCN. This mesoscale assembly accounts for parallel fibres that cross the cerebellum on the transverse axis^42,43^. Moreover, it accounts for inhibitory projections from PC to DCN ^44^ and for feedback projections from DCN to the cerebellar granule layer^18,45–47^. These connections set up intra-cerebellar communication allowing information to be transmitted in feed-forward and feedback loops. This organization is instrumental to generate cerebellar cortex activity of GrCs, GoC, MLIs, and PCs in the physiological frequency range along with the emergence of rhythmic oscillations and coherence in the circuit^47,48^. The identity of different cerebellar regions was ensured by the independence of extra-cerebellar inputs. In summary we observed the following:

1. Average discharge frequency. The oscillation frequency of cerebellar neurons in the multi-node CRBL MF, both in open and in closed loop, was compatible with that observed experimentally^32^. This ensured that the MF model was working within its natural operating regime.
2. Oscillations. Neuronal discharge showed rhythmic components on the main frequency bands revealed by MEG and EEG recordings in humans with a prevalence of the theta band^37^. This is an important aspect of validation since the CRBL MF, although tuned on mice data, proved able to capture ensemble properties typical of the human brain ^26^.
3. Coherence. Oscillations were synchronized across cerebellar regions revealing a coherence similar to empirical recordings. The cerebellar connectivity and specific neuronal dynamics of the multi-node CRBL MF support the resting-state cerebellar network revealed by fMRI recordings.

The CRBL MF model allowed the analysis and interpretation of physiological signals and their correlation with specific neuronal populations. First, the BOLD signal of the cerebellum can be directly computed from GrC activity. GrC clusters undergo sharp on-off transitions that are capable of controlling neurovascular coupling through nitric oxide (NO) production and diffusion both in the granular and molecular layer^40^. Conversely, PC activity is almost stationary over the integration time of fMRI. This aspect can impact on resting-state measurements (like here), but it would become even more relevant in task dependent fMRI simulations. Secondly, the PSD can be directly related to PCs and MLIs. PCs and MLIs, given their elongated dendritic arborization and ordered spatial orientation, generate the signals revealed by MEG and by (the rare) hd-EEG recordings available for the cerebellum^49^. Conversely, GrCs are organized in closed-field and are unlikely to contribute to these electrical signals remarkably. Therefore, signals recorded from the cerebellum can be connected to the activity of specific cell populations that are more likely to generate them.

### 3.2 The reconstruction and personalization of intra-cerebellar connectivity

A crucial step to generate a microcircuit-specific MF model of the CRBL was the curation of the intra-cerebellar SC. Even though tractography represents the gold-standard for computing the *in-vivo* human SC and pfs have been observed with histological MRI^42,50^, detection of pfs pathways *in-vivo* is impractical due to their sub-millimeter size and the specific location in the outermost layer of the cerebellar cortex. Moreover, although MRI tractography may reveal U-fibres connecting adjacent lobules, these are spurious reconstructions of under-resolved bifurcating mossy fibres. Therefore, MRI tractography alone would result in false negative and false positive connections between adjacent regions of the cerebellar cortex. Here, we first anticipated the general network connectivity and then inferred the connection weights by implementing an across-scale procedure. At a microscale level, region-specific microcircuit representations of the cerebellar network give the unique opportunity to extract synaptic convergences of pfs (see section 2.4.2) for each connection projecting from GrC of one node to GoC, MLI, and PC of transversally connected nodes (e.g., from lobule VI right/left to vermis V). This microscale reconstruction reflects a general circuit organization that needs to be scaled for the single subjects. To do so, the macroscale connectivity strength between interconnected nodes was calculated by weighting synaptic convergence values by the corresponding cerebellar volumes. In this way, the microscopic organization of the cerebellar cortical circuit was maintained and made subject-specific. A similar reasoning was applied to connections between the cerebellar cortex and DCN. The corresponding axons travelling from PC to DCN and from DCN to the cerebellar granule layer cannot be detected by MRI tractography. Again, the synaptic connectivity between cerebellar cortex and DCN was taken from microscale representations, using convergence weights that were then scaled for the corresponding regional cerebellar volumes in single subjects. To implement the inhibitory action that PCs exert on DCN, though, the sign of cerebellar cortex – DCN communication was inverted. It should be noted that CRBL MF in cortical nodes and WW models in DCN naturally communicate through spike frequencies. Thus, the cMF-TVB incorporating the multi-node CRBL MF effectively solves the major anatomical and physiological issues of signal communication in the cerebro-cerebellar loop.

### 3.3 Emergent brain rhythms in CMF-TVB simulations

Rhythmicity and multi-node coherence in cerebellar signals emerged on multiple bands during resting-state cMF-TVB simulations, in line with experimental recordings. All MF elements (i.e., GrC, GoC, PC, MLI) showed oscillations on multiple bands, although with some differences. It should be noted that oscillations in a MF element imply synchronization of the entire neuronal population in the corresponding spiking neuronal network.

The theta band was observed both in the granular and molecular layer with a dominant frequency at 6 Hz, which corresponds to the resonance frequency measured in granule cells and Golgi cells as well as in the whole granular layer^38,51,52^. In the MF, theta band oscillations are an emergent property of the cerebellar model in response to inputs from the rest of the brain. Interestingly, theta band oscillations have been revealed during control of continuous movements operated by the cerebellar-thalamic-cortical loop^37^. Moreover, theta band oscillations are associated with memory, opening new perspective to investigate the cerebellar role in memory encoding and retrieval^47,53–55^.

The alpha band was more visible in GoC and MLI with a dominant frequency at 11 Hz. This is around the spontaneous oscillation frequency of these neurons and is characteristic of the µ-wave associated with sensorimotor activity. The cMF-TVB can thus detect a fundamental correlate of the role of the cerebellum-DCN in motor timing, which indeed occurs around 10 Hz^37,56^

The beta band emerged in all cell populations, with a higher contribution in PCs, with primary frequency of 20 Hz, demonstrated the capability of cMF-TVB to detect higher rhythms emerging in both motor and cognitive functions^57,58^. Although cMF-TVB simulations are at resting state, the capability of detection of beta band can be used to understand the neuronal mechanisms underlining complex cognitive task such as semantic prediction^54^.

The gamma band shows the lowest power in MF elements but the highest PSD value, which is centered at 64 Hz, suggesting that the maximum degree of cerebellar activity synchronization occurs around the middle of the gamma band. The combination of theta and gamma waves is critical for cognitive processing, in which the cerebellum is now known to be involved^59,60^.

Abnormal cerebellar rhythms are implicated in several neurodegenerative and psychiatric conditions. In Parkinson’s disease, reduced alpha band correlates with motor impairment, while altered gamma band synchronizations correlates to the severity of tremor^61–63^. In Parkinson’s disease, a critical role of the cerebellum in cortical beta band oscillations has been suggested^64^. Furthermore, gamma and theta band alterations are observed also in schizophrenia patients^65,66^. In dementia and encephalopathy there are also distinctive EEG signatures (https://emedicine.medscape.com/article/1138235-overview?form=fpf). Therefore, the cMF-TVB embedding multi-node CRBL MF could be used to investigate the physiology and pathophysiology of cortico-cerebellar interactions in resting-state simulations as well as in sensorimotor and cognitive tasks.

### 3.4 Considerations on the performance of simulations using the multi-node CRBL MF

The multi-node CRBL MF surpassed the corresponding models using neural masses in BOLD fMRI simulations, determining a ∼50% MAE reduction compared to WW, WC, and GO (both on average and in single subjects). It should be noted that, with the multi-node CRBL MF either in open or in closed loop, neuronal populations operated within their physiological activity range with a dynamical heterogeneity driven by the specific node connectivity. Thus, the maintenance of cerebellar biological properties of neurons and microcircuit connectivity impacted favorably on the resolution of local circuit activity. Interestingly, the CRBL MF also impacted on connected regions, reducing the MAE in DCN and in the whole brain, although nodes outside the cerebellum were modeled with unspecific WW neural masses. This demonstrates the remarkable impact of integrating even a single region-specific model into virtual brain models on local and whole brain dynamics.

Since TVB is data-driven, the quality of SC is crucial both for the accuracy and the interpretation of simulations. Thus, any improvements in MRI data or the combination with other datasets (e.g., EEG or MEG) is likely to improve the simulations. We have introduced two elements of novelty: curation of the intra-cerebellar SC using a general architecture derived from anatomy *a priori* and subject-specific connection weighting using morphological parameters extracted from standard 3DT1 images. This new procedure, which resolves the inability of MRI in detecting details of local network connectivity but still allows for personalization, could be easily applied to any local networks improving the digital twin technology. Further improvements are expected from the application of ultra-high field DWI methods, which could enhance the characterization of intra-cerebellar connectivity.

### 3.5 Conclusions

This work shows that integrating anatomo-physiological properties of a specific brain circuit into large scale simulations can advance current technology toward the generation of effective brain digital twins^7^. The use of MF models to substitute specific regional nodes is promising as it improves model performance toward a closer matching with subject data. Moreover, the cMF-TVB allows to identify physiological properties of the circuit, like rhythmic oscillations and coherence, and to improve the identification of region-specific activities. The mechanisms of circuit activity could be directly related to specific neuronal populations further improving the interpretation of electrophysiological and MRI signals. Among possible developments, the reconstruction process reported here for the multi-mode CRBL MF and cMF-TVB could be applied to pathological conditions, in which patient’s circuits could be modelled according both to pathology-specific and subject-specific abnormalities. And by integrating others region-specific MFs^25,28,67,68^ one can envisage the generation of an all-MF virtual brain that could be used both for resting-state and task-dependent fMRI analysis. Eventually, cMF-TVB models embedding MF of specific brain regions could be used for the investigation of sensorimotor and cognitive processes and to generate virtual twins of patient’s brain in pathological conditions, opening new opportunities for diagnosis and treatment.

## 4. Material and Methods

TVB simulates subject-specific brain dynamics at different scales starting from SC matrices, usually derived applying tractography to diffusion weighted imaging (DWI) data^2^. TVB allows simulating the neuronal activity of interconnected regions by resolving computational models of local circuit function in each node. By convolving the resulting neuronal activity with a built-in hemodynamic response function, TVB provides a simulation of the BOLD signal for each node^3^. Applications of TVB in clinical trials and research studies are growing, and include the improvement of pre-operative planning in epilepsy and the investigation of subject-specific metabolic changes in neurodegenerative diseases^4,11–17^.

MRI data was downloaded from the Human Connectome Project (http://db.humanconnectome.org). Cerebral and cerebellar nodes were identified based on an ad-hoc atlas and were used to compute the empBOLD time-series, from resting state fMRI data (section 4.1), and the whole brain SC, from DWI data.

The workflow is reported in supplementary material (Fig. S1) and all the steps are explained in detail in sections 4.2 to 4.7. Briefly:

A. Curation of the cerebellar SC to account for cerebellar-specific connectivity (section 4.2). We differentiated pfs axonal pathways, connecting adjacent cerebellar cortex regions, from the mfs pathways, connecting the cerebellar cortex with DCN and/or with the cerebrum.
B. Integration of CRBL MF equations in TVB (sections 4.4-4.5). We associated one CRBL MF to each node of the cerebellar SC, and a generic model to DCNs and cerebral nodes. We updated the model equations to simulate the propagation of the neuronal activity from one region to another one, and we updated the input/output relations to enable generic models (e.g., WW) receiving inhibitory input coming from the CRBL MF.
C. TVB simulations (section 4.6). We tested the framework in two conditions: first, the cerebellum in open-loop (i.e., isolated cerebellar cortex activity) to assess the performance of the *multi-node CRBL MF*; then, the cerebellum in closed-loop (i.e. whole brain activity) to assess the performance of the *cMF-TVB* including different models for different regions.
D. Validation (section 4.7). Multi-node CRBL-MF and cMF-TVB simulations were evaluated in terms of constructive and predictive validity.

### 4.1 Dataset and data preparation

High-quality pre-processed MRI data (Siemens 3T Connectome Skyra MRI scanner with a 32-channel receive head coil) was downloaded from the Human Connectome Project database (https://db.humanconnectome.org/) ^69^. The dataset included the following imaging data: DWI (1.25 mm isotropic resolution, b = 1000, 2000, 3000 s/mm^2^, 90 isotropically distributed directions/b-value and 18 b_0_ images), rs-fMRI (2 mm isotropic resolution, TR/TE = 720/33.1 ms, 1200 volumes) and 3DT1-weighted images (0.7 mm isotropic resolution). We selected the same 8 healthy subjects (2 males and 6 females; 30.6 ± 4.1 years) used in our previous work on the impact of cerebellar connectivity on whole brain dynamic simulations^10^.

#### 4.1.1 Brain parcellation

Brain regions were identified using an already defined ad-hoc atlas including cerebellar parcellations from a spatially unbiased atlas template of the cerebellum and brainstem (SUIT), cerebral parcellations from automated anatomical labeling (AAL) atlas, and deep gray matter structures using FSL-first ^10^. Whole brain parcellations resulted in 126 regions with 27 cerebellar cortical regions, 6 deep cerebellar nuclei, and 93 cerebral regions. Each gray matter parcellation was considered as a node for the SC extraction and for the assignment of a computational model to run TVB.

### 4.2 Structural Connectivity (SC)

For each subject the SC matrix was computed using the anatomically constrained tractography with 30 millions of streamlines implemented in MRtrix3^70,71^. SC weights were defined using probabilistic streamline tractography (second-order integration over fiber orientation distributions – (iFOD2)) and by assigning each streamline to the nearest node in a 2-mm spheric neighborhood^72^. Elimination of spurious ipsilateral cerebro-cerebellar tracts and the selection of contralateral efferent and afferent cerebellar connections were performed from whole brain tractograms as described in^29,30,41^. Directionality of the cerebro-cerebellar loop connections were included in the SC matrix by imposing the assumption that cerebellar efferent streamlines are via the superior cerebellar peduncle while the cerebellar afferent streamlines are via the middle cerebellar peduncle^10^.

#### 4.2.1 Curation of the intra-cerebellar SC

The cerebellar cortex SC was curated to introduce the missing connectivity between adjacent cerebellar cortical regions, which cannot be detected from DWI tractography. Therefore, we introduced weights driven by known properties of pfs (Fig. 2A). Moreover, to provide a realistic connectivity weight for the cerebellar cortex-DCN loop, we used known properties of PC axonal bundles (Fig. 2B)^73,74^. These steps are provided in detail below:

##### Cerebellar cortex connectivity

Connectivity weights between pair of adjacent regions, driven by pfs, were quantified with a multi-scale approach by extracting the pfs synaptic convergence from a previously validated cerebellar cortex SNN and weighting it with subject-specific morphological constraints^33,34,42^. Specifically, the synaptic convergence values from GrCs on GoCs, on MLIs and on PCs were weighted by the volume of each pair of interconnected cerebellar regions to account for inter-subjects’ variability, and then normalized for the intra-cranial volume to prevent any bias. Volumes of cerebellar regions were computed by overlapping the SUIT atlas with the anatomical 3DT1. The SUIT atlas provides separate parcellations for the Vermis and the right and left lobules, thus the topology of the cerebellar cortex SC was defined for adjacent regions k (with k from I to X): lobule(k) left – vermis(k) and lobule(k) right – vermis(k) (e.g., lobule IX right – vermis IX and lobule IX left – vermis IX; see supplementary material – Fig. S2).

##### Cerebellar cortex – deep cerebellar nuclei connectivity

Connectivity weights from cerebellar cortex regions to DCN, driven by PC axons were quantified as explained in 4.2.2, and assigned to the efferent (i.e., from cerebellar cortex to DCN) bundles. Specifically, a streamline was assigned to all the nodes encountered along its whole length^72^. The weight of afferent bundles (i.e., from DCN to cerebellar cortex) was computed as the 10% of the correspondent forward projections^74^. In order to correctly link cerebellar cortex with other brain regions, it is important to make a functional consideration: while models of cortical regions are connected to each other through excitatory fibres, similarly to what happens between two adjacent CRBL MF transmitting excitatory signals, each CRBL MF has an inhibitory connection to the DCN^44,75^. As in TVB the connectivity between functional models is weighted by the corresponding SC, to capture the inhibitory/excitatory nature of the signals we multiplied all connections from the cerebellar cortex to the DCN by the value-1.

#### 4.2.3 Functional Data

The time-course of the BOLD signal was extracted from resting-state fMRI for each region, resulting in a 14-minute signal, namely the empBOLD signal. The empBOLD was used as ground-truth to evaluate the simBOLD signals (section 4.7)

### 4.4 Multi-node CRBL MF

A multi-node CRBL MF was implemented by associating a CRBL MF model to each node of the cerebellar cortex to test the CRBL MF integrated into the TVB neuroinformatic platform in an open loop configuration.

The CRBL MF was optimized as follows, considering that for intra-cerebellar cortical connections the input of a CRBL MF needs to be driven by the pfs, while the background noise mimicking the input from the cortex and DCN is driven by mfs

#### 4.4.1 Transfer Function modifications

Population-specific Transfer Functions (F in equations) of GoC, MLI and PC integrated also the input carried by the pfs of the GrC to connected CRBL MF.

As an example, from the single-node CRBL MF^26^ to the multi-node CRBL MF, the PC transfer function was updated from:

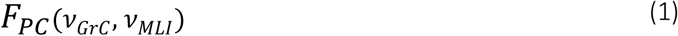

to

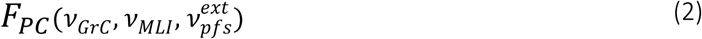

where *ν*_GrC_ is the activity of the GrC of the same module, *ν*_MLI_ the activity of the Molecular Layer Interneurons of the same node, and *ν*_pfs_^ext^ is the external activity with respect to the module, i.e., the activity of the GrC population of the adjacent modules driven by parallel fibres.

Following the same rationale MLI Transfer Function and GoC Transfer Function resulted in:

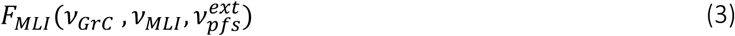

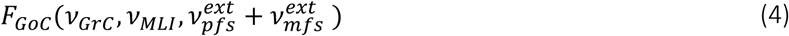

The GrC transfer function was invariant since the GrC population receives an external driven only from the mfs:

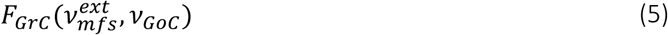

#### 4.4.2 Input configuration

In TVB the total input *c* ([Hz]) for a target node *k* is computed as the summation of the activity of all the afferent nodes weighted by the SC and resulting in:

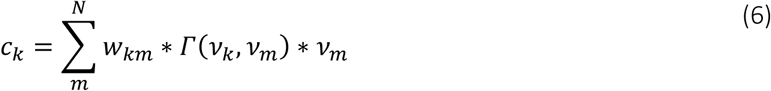

where *N* is the total number of nodes, w_km_ is the SC weight from node *m* to the target node *k*, Γ is the coupling function between node *m* and target node *k* and *ν_k_*, *ν_m_*, are the activity of node *k* and node *m*, respectively. Activity of the target node *m* at the time instant *t* is computed by summing the activity of the target node recorded at the previous time instant *τ = t-dt* (dt = integration step) and the input *c* from afferent nodes:

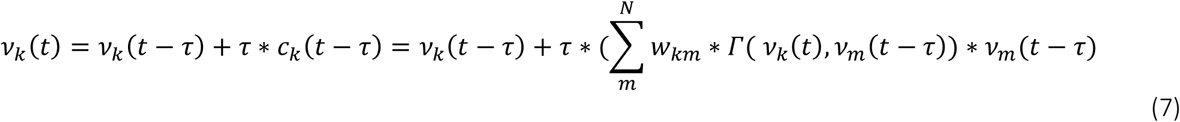

To differentiate the contribution of mfs and pfs, c was weighted for the mfs convergence and the pfs convergence estimated from a previously validated SNN used to construct the CRBL MF^26,31,33,34^ resulting in cMF and c_pf_ (Table 4). To account for the convergence heterogeneity across cerebellar neuronal populations (e.g., GrC receives only cMF, while GoC both cMF and c_pf_), cMF and c_pf_ were further split into contributes specific for each population as reported in Table 5.

**Table 4.**
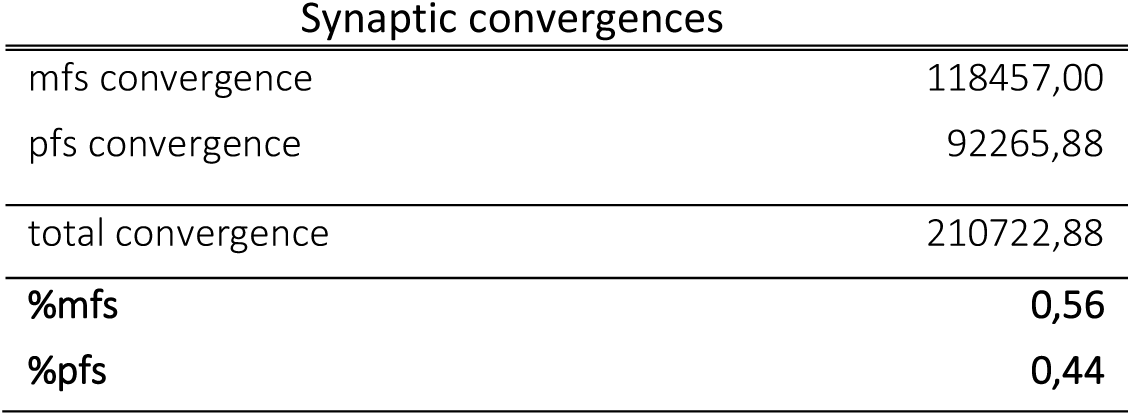
Synaptic convergence and divergence. Synaptic convergence of mfs and pfs extracted from the cerebellar spiking neural network. Total synaptic convergence was computed by summing mf and pf synaptic convergence and the rate of mfs and pfs was computed on the total synaptic convergence. The mf ratio (%mfs) was computed as the ratio between mf convergence and total convergence. The same rationale was applied to compute the pf ratio (%pfs). The total input (c [Hz]) to a node i, was differentiated from input carried by mfs and input carried by pfs as: c_mfs_ = c_i_ * mfs% and c_pfs_ = c_i_ * pfs%.

**Table 5.** Circuit connectivity. Mfs and pfs contribution, specific for each cerebellar neuronal population computed as C_mfs_^P^ = C_mfs_ *%mfs^p^ and C_pfs_^p^ = * %pfs^p^, with p = neuronal populations receiving input c In open loop (i.e., isolate cerebellar cortex), GrCs of different CRBL MF are not connected (c_pfs_^GrC^ = 0) while MLIs and PCs receive input only through GrCs of interconnected models (c_pfs_^MLI^ ¹ 0 and c_pfs_^MLI^ ¹ 0). According to the construction of the CRBL MF, GoC is the only population with a double-input channel (c_pfs_^GoC^ ¹ 0 and c_mfs_^GoC^ ¹ 0).

| Incoming connection (To) | $c_{mfs}^p$ | $c_{pfs}^p$ |
| --- | --- | --- |
| GrC | 0,97 | 0 |
| GoC | 0,03 | 0,14 |
| MLI | 0 | 0,55 |
| PC | 0 | 0,31 |

This estimation was necessary in the open loop configuration only, because the external input was superimposed to all the nodes as a background noise mimicking the input from the cerebrum and/or the DCN. In the closed loop scenario, this superimposition was not required because the external input from cerebrum and/or the DCN was simulated.

### 4.5 Construction of the cMF-TVB

The cMF-TVB was implemented considering the cerebellum wired in a closed loop with the DCN and the cerebrum. The CMF-TVB was built by interconnecting 126 models (one per region), divided into 27 CRBL MFs, corresponding to the cerebellar cortex, and 99 WW models, corresponding to the cerebral cortical and subcortical regions, as well as the DCN^20,26^. A schema of the inter-models connectivity is reported in Fig. 3. The cMF-TVB inherits the population-specific transfer functions, and the input configurations described in 4.4.1 and 4.4.2 for the multi-node CRBL MF. In addition, it requires further modifications to the input/output relations of the CRBL MF (4.5.1) and the WW models (4.5.2) to allow for inter-models connections.

#### 4.5.1 Cerebellar mean field model (CRBL MF)

Cerebellar MF equations were modified in terms of population specific transfer function to include the coupling contribution following the same rationale detailed in 4.4. With this configuration, the contribution carried by pfs (i.e., from adjacent cerebellar MFs) was separated by the one carried by other region nodes (i.e., from a connected WW), resulting in a physiological mapping of the input/output relations. Indeed, the input carried by pfs projecting from GrCs of adjacent nodes became the part of the input to GoC, MLI and PC of the target module, while the granular layer of the target module was driven by the input from mossy fibres from the connected DCN and/or cerebral models.

In detail, with this configuration, each population of a CRBL MF could receive an additive “external input” coming from adjacent modules. The coupling is specifically set up according to the origin of the external input: in the case it comes from a cerebellar cortex module to another cerebellar cortex module it goes from the GrC of the source module to the GoC, MLI and PC of the target module; on the other hand, in the case it comes from a DCN or cerebral node (e.g., from a WW model) the input goes to the granular layer of the target cerebellar module, namely to the GrC and GoC. Therefore, in a mathematical form, the coupling towards any cerebellar cortex node can be written as:

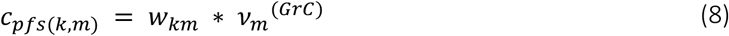

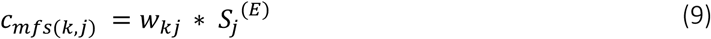

where k is the cerebellar target node (e.g., vermis X), w_km_ is the curated SC weight from cerebellar source node m (e.g., right lobule X), v_m_^(GrC)^ is the activity of the GrC population in node m, w_kj_ is the SC weight that can be either from a cerebral node or a DCN node (e.g., right fastigial), and S_j_^(E)^ is the activity coming from the WW excitatory population of such cerebral or DCN node.

It is worth noting that within the cMF-TVB, the differentiation between mossy and parallel fibres is achieved by segregating the input activity to a cerebellar cortex target node, which is driven separately by parallel and mossy fibres. Consequently, the estimation of mossy fiber parameters and coupling strength (section 4.4.2) is not required.

#### 4.5.2 Wong Wang model (WW)

The excitatory population of the WW model (E) receives input both from other Es belonging to other modules of connected WWs, and from PCs of connected cerebellar MFs. Therefore, it was necessary to modify the WW equation to separate the excitatory contribution from Es and the inhibitory contribution of PCs. The WW coupling expression (coupling_j_) for a target node j (e.g., the right fastigial), was modified from Deco et al., 2014^20^ as follows, by adding the inhibitory input from PCs:

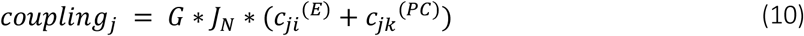

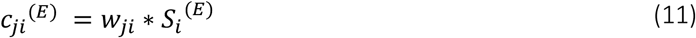

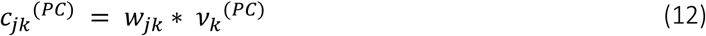

Where G is the global coupling parameter, J_N_ is the synaptic parameter associated with the NMDA channel, c_ji_^(E)^ is the excitatory activity of cerebral source node i (S_i_^(E)^), weighted by the structural weight w_ji_ between target node j and source node j. c_jk_^(PC)^ is the PC activity from the cerebellar source node m (v_k_^(PC)^) weighted by the structural weights w_jk_ to target node j from source node k (e.g., to right fastigial nucleus from Vermis X, see Fig. 3). Note that the indicial notation subtends a summation over index i and k.

With the addition of c_jk_^(PC)^, it is possible to introduce the contribution of a cerebellar-specific inhibitory activity into the WW equations^20^. Both external excitatory and inhibitory input for a target node k (I_j_^(E)^ and I_j_^(I)^ respectively) are dependent on specific coupling terms (c_jk_^(E)^ and c_ji_^(PC)^):

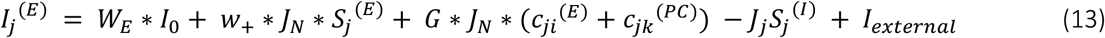

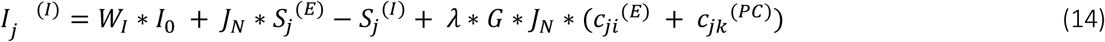

Where W_E_ and W_i_ are scaling parameters of the overall effective external input (I_0_ = 0.382 nA), w+ is the local recurrent excitation, J_N_ is the parameter associated to NMDA channel, S denotes the average synaptic gating variable of target node k, J_i_ is local inhibitory current, I_external_ is the external stimulation for simulating task evoked activity (set at 0 for resting-state simulation), and λ indicates the long-range feedforward inhibition. The superscripts (E), (I) and (PC) indicate the populations as introduced above.

The coupling term for the cerebellar nodes was differentiated into contribution from parallel and contribution from mossy fibres (c_pfs(k-m)_ and cMF_s(k-j)_ respectively), resulting in the following expression for a cerebellar target node k (e.g. Vermis X), receiving input from a cerebellar source node m (e.g., right lobule X), from a DCN source node j (e.g., right fastigial nucleus), and from a cerebral node (e.g., thalamus):

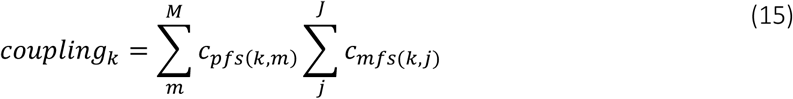

Where m =cerebellar cortical node, and j = DCN and/or cerebral node, and c_pfs(k-m)_ and cMF_s(k-j)_ defined as in section 4.5.1

### 4.6 Simulations

Simulations lasting 14 minutes, with an integration step (dt) of 0.1 ms, were performed to investigate cerebellar activity in both open-loop and closed-loop configurations. In the closed-loop simulations, the impact of cerebellar activity on whole brain dynamics was also analyzed. Neuronal activity and BOLD signal were simulated for each node following the set-up reported in Table 6.

**Table 6.** Simulation parameters. Configuration of the simulation parameters used to test both the multi-node cerebellar mean field and the hybrid virtual brain. The set-up of these parameters is required to run simulations using TVB neuroinformatic platform.

| Simulation set-up |  |
| --- | --- |
| Simulation length | 14 min |
| Neuronal activity integration step | 0.1 ms |
| Neuronal activity transient | 1/3 of simulation length |
| BOLD signal integration step (TR) | 720 ms |
| BOLD transient | 36 s |
| Integrator | Heun stochastic |
| Integration noise | Additive random |

#### 4.6.1 Multi-node CRBL MF simulations (open loop)

The cerebellar cortex SC was used to map the connections strength between CRBL MFs constructing the multi-node CRBL MF. Each node of the cerebellar cortex SC was associated with a model, resulting in a network of 27 interconnected models. The cerebellar cortex activity was simulated comparing the performance of generic models already available in TVB, namely Wong Wang model (WW), Wilson Cowan model (WC), and Generic Oscillator (GO) against the performance of the region-specific CRBL MF, purposely integrated in TVB.

#### 4.6.2 cMF-TVB simulations (closed loop)

Whole brain SC was used to map the connections strength between models in cMF-TVB. Whole brain activity was simulated either with a standard virtual brain (TVB), using the WW model for all nodes, or with the cMF-TVB, combining 93 WW models covering the cerebrum and the DCN with 27 CRBL MF models for the cerebellar cortex.

### 4.7 Validation

Multi-node CRBL MF and cMF-TVB simulations were evaluated both in term of constructive and predictive validity. Moreover, cMF-TVB simulations were used to explore the cerebellar emergent brain rhythm, assessing the contribution of each population to specific frequency bands, and their propagation over the cerebella regions. Finally, emergent synchrony amongst the cerebellar nodes was computed in term of coherence.

#### 4.7.1 Constructive validity

Simulated cerebellar neuronal responses were compared with the physiological ranges of each neuronal population both in open and in closed loop to assess whether the constructive validity, already demonstrated for the CRBL MF model itself, was maintained in both multi-node CRBL MFs framework and cMF-TVB.

#### 4.7.2 Predictive validity

The TVB platform was applied to simulate BOLD signals using neuronal models as generative models. The ability of the cMF-TVB to simulate BOLD signals using regions-specific models as generative sources was quantified by computing the MAE between the empBOLD and the simBOLD signal. We computed the overall MAE and the subject-specific MAE.

Shapiro-Wilk and Levene’s tests were applied to check the normal distribution of the overall MAE for each model (i.e., CRBL MF, WW, WC, and GO), and the homogeneity of variance across models. Based on the outcome of the Shapiro-Wilk and the Levene’s test, Mann-Whitney was selected as the most appropriate statistical test to compare the distributions of the overall MAE between pair of models. The statistical significance threshold was set to 0.001.

#### 4.7.3 Emergent cerebellar rhythm: a frequency band exploration

Power Spectrum Analysis was implemented to investigate the predominant frequency band of the activity of each cerebellar neuronal population and, consequently, the overall cerebellar dominant frequency spectrum. For each subject, the cerebellar population-specific Power Spectrum Density (PSD) was computed with the Fourier transform, using the Welch method to reduce the PSD variance. Sampling frequency was set to 250 Hz. PSD was analyzed firstly (i) considering the cerebellum as a unique region, and then (ii) considering the cerebellar cortical regions separately:

i. For each brain rhythm band, PSD was averaged over the cerebellar regions. The Area Under the Curve (AUC) of each population-specific PSD was computed and normalized for the number of frequencies in each band (AUC_norm_), then it was averaged across subjects to quantify the total predominant activity band for each population. A band-specific predominant frequency was then averaged across all the cellular populations to identify the cerebellar peak frequency within each band.
ii. For each brain rhythm, for each region, AUCs_norm_ were computed to explore the frequency bands propagation. The rate of brain rhythm bands within each region was obtained by summing the population-specific AUC_norm_, so providing the overall brain rhythm band distributions across cerebellar regions.

#### 4.7.4 Synchrony in the cerebellar subnetwork

For each subject, the coherence was computed on the CMF-TVB simBOLD, on the TVB simBOLD and on the empBOLD of the cerebellar cortex. For each pair of cerebellar cortical nodes, the coherence was computed as:

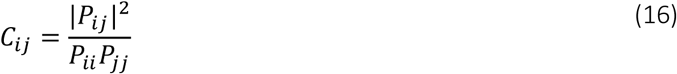

Where C is the coherence between node i and node j, P_ij_ is the cross-spectral density estimate from the BOLD signal recorded from x and y, P_ii_ and P_jj_ are the PSD of the BOLD signals of node i and j, respectively. PSDs were computed by applying the Fourier transform to the BOLD signals divided in windows of length = 512, with an overlap of half window length (i.e., 512/2) and a sampling frequency as the TR (0.72 s).

For each subject, coherence matrices for cMF-TVB, TVB and empirical fMRI data were computed by averaging the coherence resulting for each frequency in the spectrum.

To quantify the discrepancy between simulated and empirical data, cMF-TVB and TVB coherence matrices were compared to that resulting from the empirical BOLD using the following scores: (i) MAE to provide a measure of how close the matrices are in terms of coherence strength, (ii) Frobenius Norm to quantify the overall deviation between the two matrices, (iii) cosine similarity to capture the overall difference in the orientation of the two matrices, (iv) Kullback-Leibler (KL) divergence to compare the distribution of the two matrices, (v) Pearson correlation coefficient (PCC) and Spearman correlation coefficient (SCC) to compare the trend of the coherence values. Significant differences were computed using a paired t-test on cMF-TVB vs empirical data and TVB vs empirical data, separately. Significance threshold was set at 0.001.

### 4.8 Pipeline availability

Our pipeline is entirely available, including the curation of the intra-cerebellar SC and the integration and interconnection of the CRBL MF. Implementation of intra-cerebellar SC curation is available at https://github.com/RobertaMLo/tvb_data_preprocessing. The cerebellar TVB is built as a python package consistent with the AdEx TVB previously implemented and it is available under requests at https://github.com/RobertaMLo/IntegrationMFintoTVB, which includes the cerebellar MF model equation (`parallel_crbl.py`) and the configuration file of cerebellar-specific parameters (`parallel_crbl_params.py`)

## Notes

### Competing Interest Statement

The authors have declared no competing interest.

