## Supplementary Material for "Region-specific mean field models enhance simulations of local and global brain dynamics"

### Supplementary Materials

The pipeline used to develop and validate the cMF-TVB is reported in Fig. S1. An insight on the approach used to implement the curation of the intra-cerebellar SC by combining macroscale information derived by MRI data and microscale SNN is reported in Fig. S2.
An example of a subject-specific PSD is shown in Fig. S3. The frequency-band AUC was computed considering the frequencies magnitude of each band separately. Table S1 reported for each region the population-specific carrier frequency resulted from the PSD averaged over the subjects.

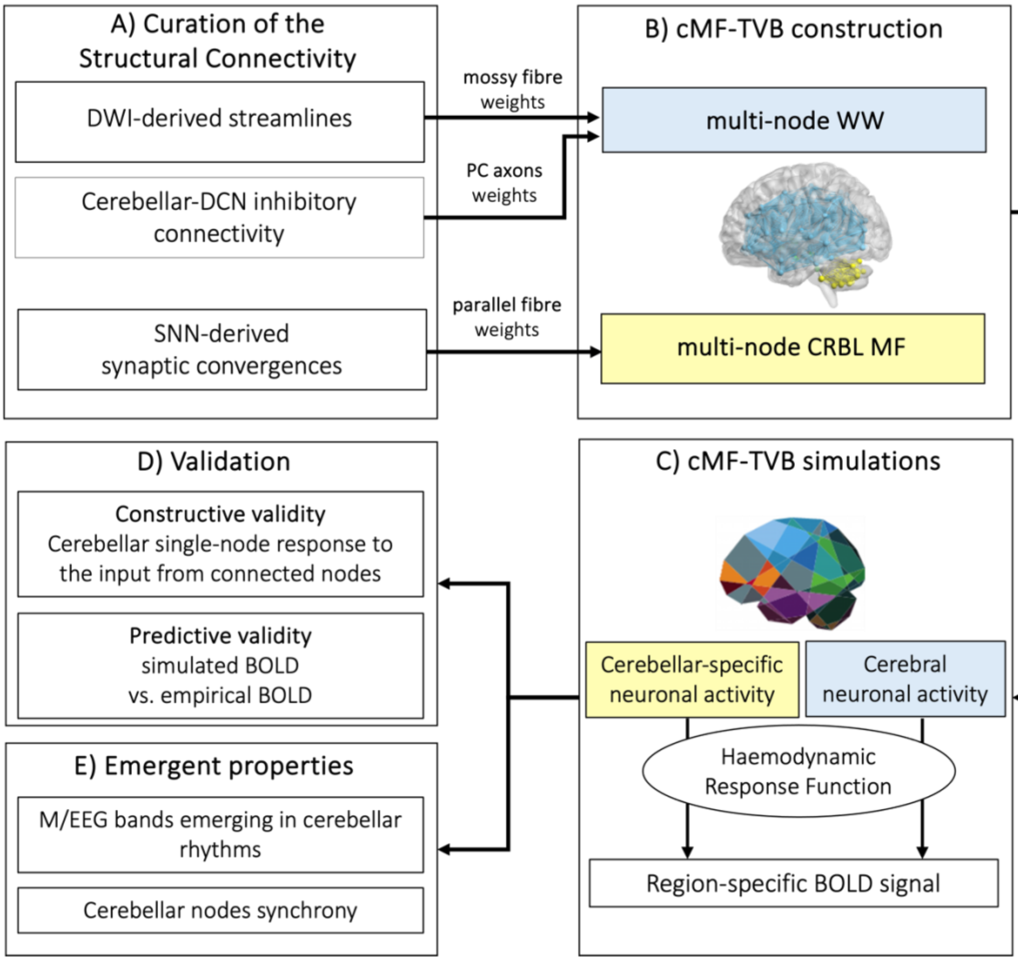


**Fig. S1) Detailed technical pipeline. A)** Curation of structural connectivity involves presetting cerebellar-DCN network wiring and then weighting it for the specific subject anatomy. **B)** The standard oscillator, i.e., WW model, was associated to cerebrum and DCN nodes, while the CRBL MF to the cerebellar cortex nodes. **C)** TVB simulations were performed simulating firing rate and BOLD signals for each node. **D)** cMF-TVB were evaluated in terms of their constructive and predictive validity. **E)** Emergent properties were evaluated to assess the capability of cMF-TVB of reproducing emergent cerebellar rhythm in M/EEG and the synchronization across cerebellar regions. The framework was firstly evaluated with the cerebellum in open loop (isolated cerebellar cortex) to test the performance of a multi-node CRBL MF model and then with the cerebellum wired in a closed loop (whole-brain) to test hybrid networks.

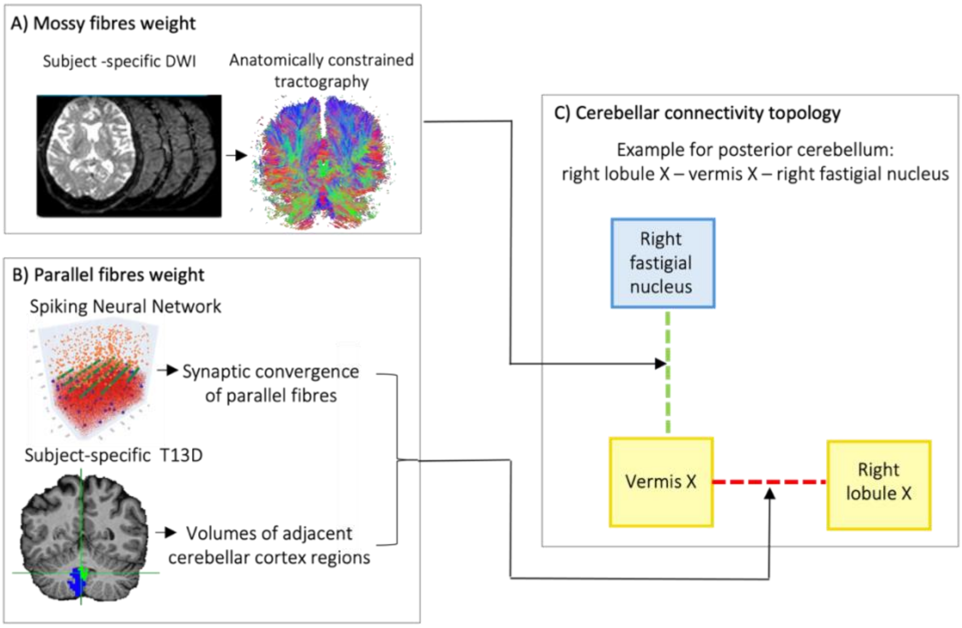


**Fig. S2) Multiscale approach for the curation of the intra-cerebellar structural connectivity. A)** The weights of mossy fibers (mfs) from DCN and/or cerebrum are computed with anatomically constrained tractography. **B)** The weights of parallel fibers (pfs) are extracted from the cerebellar spiking neural network as populations synaptic convergence and weighted with volumes of connected lobes. **C)** Cerebellar cortex-to-DCN connections are quantified as in **(A)** (green connection), intra-cerebellar cortex connections as in **(B)** (red connection).

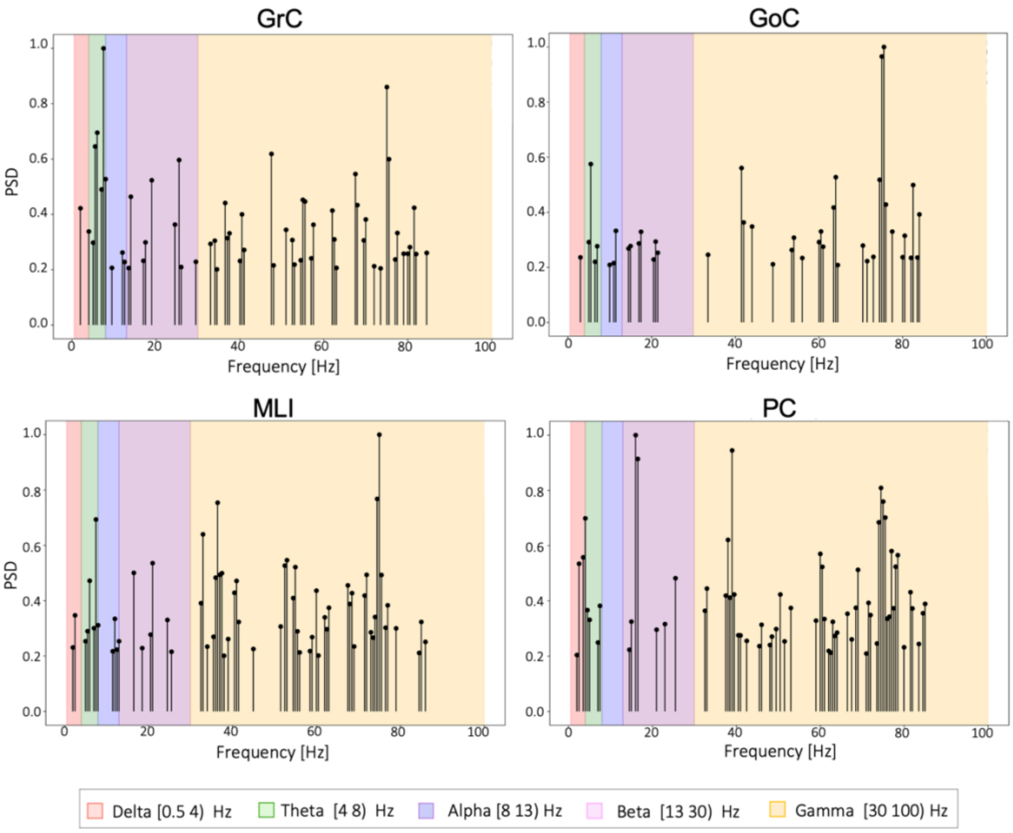


**Fig. S3) Population-specific PSD for a randomly chosen subject**.

PSD was computed separately for each population (Granule Cells (GrC), Golgi Cells (GoC), Molecular Layer Interneurons (MLI) and Purkinje Cells (PC)) by averaging populations activity on the cerebellar regions included in the structural connectivity. Brain rhythm bands are defined as: Delta = [0.4 4) Hz (red), Theta = [4 8) Hz (green), Alpha = [8 13) Hz (lilac), Beta = [12 30) Hz (purple), Gamma = [30 100) Hz (yellow). The same analysis was repeated for each subject producing subject-specific PSD.

| | Overall carrier frequency [Hz] | | | |
|---|---|---|---|---|
| Regions | GrC | GoC | MLI | PC |
| Left_I_IV | 2.00 | 81.50 | 6.50 | 18.50 |
| Left_V | 56.00 | 71.00 | 49.00 | 2.50 |
| Left_VI | 23.50 | 81.50 | 23.50 | 23.50 |
| Left_CrusI | 22.00 | 18.00 | 22.00 | 42.50 |
| Left_CrusII | 36.00 | 27.50 | 54.50 | 45.50 |
| Left_VIIb | 57.50 | 23.00 | 13.50 | 13.50 |
| Left_VIIIa | 59.50 | 37.50 | 5.00 | 59.50 |
| Left_VIIIb | 32.50 | 32.00 | 23.50 | 47.00 |
| Left_IX | 46.50 | 48.50 | 46.50 | 46.50 |
| Left_X | 61.50 | 67.50 | 53.50 | 29.00 |
| Vermis_VI | 61.50 | 79.00 | 22.00 | 22.00 |
| Vermis_CrusII | 51.00 | 31.00 | 84.50 | 59.00 |
| Vermis_VIIb | 10.00 | 39.00 | 10.00 | 48.50 |
| Vermis_VIIIa | 45.50 | 74.50 | 41.00 | 41.00 |
| Vermis_VIIIb | 22.00 | 47.50 | 84.00 | 80.00 |
| Vermis_IX | 40.50 | 62.00 | 6.00 | 54.50 |
| Vermis_X | 50.00 | 64.50 | 4.00 | 27.00 |
| Right_I_IV | 39.00 | 44.00 | 39.50 | 50.50 |
| Right_V | 3.50 | 64.50 | 3.50 | 67.50 |
| Right_VI | 84.00 | 1.50 | 80.00 | 80.00 |
| Right_CrusI | 68.00 | 62.00 | 34.50 | 34.50 |
| Right_CrusII | 51.00 | 36.50 | 39.00 | 76.50 |
| Right_VIIb | 54.00 | 1.00 | 48.50 | 11.00 |
| Right_VIIIa | 3.50 | 8.50 | 3.50 | 41.00 |
| Right_VIIIb | 33.00 | 47.00 | 87.00 | 38.50 |
| Right_IX | 6.00 | 4.50 | 6.00 | 6.00; 20.50 |
| Right_X | 74.00 | 33.00 | 74.00 | 77.00 |

**Table S1. Population-specific carrier frequency.** Carrier frequency was computed as the frequency with the maximum intensity in the PSD analysis.
